# Regulatory networks of gene expression in maize (*Zea mays*) under drought stress and re-watering

**DOI:** 10.1101/361964

**Authors:** Liru Cao, Xiaomin Lu, Pengyu Zhang, Lixia Ku, Guorui Wang, Zhen Yuan, Xin Zhang, Jingyu Cui, Jingli Han, Ying Liu, Yangyong Cao, Li Wei, Tongchao Wang

**Affiliations:** Collaborative Innovation Center of Henan Grain Crops, Agricultural College of Henan Agricultural University, Zhengzhou, Henan 450046, China; National Engineering Research Centre for Wheat; Grain Crops Research Institute, Henan Academy of Agricultural Sciences, Zhengzhou 450002, China

**Keywords:** Water-deficit, water-sufficient, RNA-sequencing, drought-resistance, proline synthesis, cell wall, ABA, molecular mechanisms

## Abstract

Drought can severely limit plant growth and production. However, few studies have investigated gene expression profiles in maize during drought/re-watering. We compared drought-treated and water-sufficient maize plants by measuring their leaf relative water content, superoxide dismutase and peroxidase activities, proline content, and leaf gas exchange parameters (photosynthetic rates, stomatal conductance, and transpiration rates). We conducted RNA sequencing analyses to elucidate gene expression profiles and identify miRNAs that might be related to drought resistance. A GO enrichment analysis showed that the common DEGs (differently expressed genes) between drought-treated and control plants were involved in response to stimulus, cellular process, metabolic process, cell part, and binding and catalytic activity. Analyses of gene expression profiles revealed that 26 of the DEGs under drought encoded 10 enzymes involved in proline synthesis, suggesting that increased proline synthesis was a key part of the drought response. We also investigated cell wall-related genes and transcription factors regulating abscisic acid-dependent and -independent pathways. The expression profiles of the miRNAs miR6214-3p, miR5072-3p, zma-miR529-5p, zma-miR167e-5p, zma-miR167f-5p, and zma-miR167j-5p and their relevant targets under drought conditions were analyzed. These results provide new insights into the molecular mechanisms of drought tolerance, and may identify new targets for breeding drought-tolerant maize lines.

**Abbreviations:** leaf relative water content: RWC, superoxide dismutase activity: SOD, peroxidase activity: POD, proline content: Pro, photosynthetic rates: Pn, stomatal conductance: Cond, transpiration rates: Tr.; quantitative real-time polymerase chain reaction: qPCR; abscisic acid; ABA; polyethylene glycol :PEG; Principal component analysis :PCA; polyacrylamide gel electrophoresis :PAGE

**Highlight:** The study of physiology and molecular mechanism of maize laid a theoretical foundation for drought resistance breeding under drought stress and re-watering.

## Introduction

Maize is one of the three major food crops and it has an ancient cultivation history. Drought is one of the most important environmental factors that severely affects plant growth, yields, and crop quality on a global scale (Boyer, 1982; Tollenaar and Lee, 2002). Cultivating fine varieties of maize with enhanced drought resistance is of great practical significance to increase maize yield. Therefore, the discovery of drought-resistance genes to breed drought-resistant maize lines is one way to address the issue of drought in the long term (Monneveux et al., 2006). Now that the whole genomes of maize and other plants have been sequenced, many researchers are taking advantage of RNA-seq to study the gene expression patterns in plants under different environmental conditions or at different growth stages. Such studies have provided a great deal of important information about the regulation of gene expression and have identified candidate genes associated with specific traits such as drought tolerance.

Transcriptome analyses have been very useful to discover the functions of various genes in maize. For example, such studies have identified genes involved in the response to various abiotic stresses such as low-phosphorus (low-P), excess nitrate, ultraviolet-B radiation, oxidative stress, and drought. A transcriptome analysis of maize seedling roots under low-P stress identified key genes involved in the response to low-P (Calderon-Vazquez et al., 2008). In another study, transcriptome sequencing of maize roots subjected to nitrate stress demonstrated the co-transcriptional pattern of nitric oxide synthesis/clearance genes in root epidermal cells (Begheldo, 2011). The responses of maize leaves and ears to irradiation of varying duration was investigated using transcriptome and metabolomics analyses, which revealed that the early response in all tissues may be caused by the same signaling pathway, while the response becomes increasingly organizational-specific with longer irradiation time (Casa ti, Morrow, Fernandes, & Walbot, 2011). Analyses of the gene expression profile of the maize Rg1 mutant revealed the roles of Rg1 in dissipating reactive oxygen species (Guan et al., 2012). A microarray analysis of the gene expression profile of maize B73 under various stress conditions found that key genes may be expressed at critical sites to regulate gene expression under different stress conditions, thereby playing important roles increasing stress resistance (Fernandes, Casati, & Walbot, 2008). Studies on the gene expression profiles of maize under drought stress have indicated that drought-resistant varieties more effectively activate the expression of drought-related genes after water stress (Hayanokanashiro, Calderónvázquez, Ibarralaclette, Herreraestrella, & Simpson, 2009).

MicroRNAs (miRNAs) are a class of non-coding small molecule single-stranded RNAs encoded by endogenous genes with a length of about 16-29 nt. The precursors of miRNAs have a hairpin structure. miRNAs were first discovered in nematodes and were subsequently predicted to exist in plants such as rice, *Arabidopsis thaliana*, maize, and *Brassica napus* (Mica, Gianfranceschi, & Pè, 2006). miRNAs can regulate gene expression by degrading target RNAs at the post-transcriptional level or inhibiting translation at the translation level (Bartel, 2004; Humphreys & Preiss, 2005). These processes participate in regulating gene expression in eukaryotic cells. In plants, genes encoding miRNAs are located in intergenic regions (Voinnet, 2009) or in gene introns (Lagosquintana, Rauhut, Lendeckel, & Tuschl, 2001). Some miRNA genes are close to each other and form gene clusters, and these miRNA genes can co-transcribe to form one primary miRNA (pri-mi RNA) (Zhang et al., 2013). Many miRNAs have been verified in plants, and have been shown to be involved in diverse stress responses (Feng et al., 2015; Jia & Tang, 2010; Jones-Rhoades & Bartel, 2004). Solexa sequencing revealed that the miR528, miRl59, miR164, miR167, miR169, miR319, and miR396 families were highly expressed in the root; and the miR156, 166, 167, and 168 families were expressed in different parts and growth stages. These findings highlighted that the abundance of miRNAs varies among different tissues (Zhang et al., 2009).

The drought resistance of plants is judged as their ability to endure drought stress and rapidly resume growth after re-watering. Being able to resume growth after re-watering is relatively more important for maize yield (Chaves & Oliveira, 2004). Few studies have investigated gene expression profiles and searched for drought-resistance genes in maize after re-watering (Chen et al., 2015). Transcriptome analysis is a useful tool to analyze the regulatory networks that operate during drought treatment. Therefore, we used RNA sequencing to analyze the gene expression profile and identify miRNAs related to drought resistance in maize during and after a drought treatment. We compared the RWC (leaf relative water content), superoxide dismutase (SOD) and peroxidase (POD) activities, proline (Pro) content, and leaf gas exchange parameters between water-deficit and water-sufficient maize plants. We also detected differences in global gene expression that could be related to the drought response. Finally, we analyzed cell wall-related genes and transcription factors regulating the abscisic acid (ABA)-dependent and ABA-independent pathways. The results of this study provide new insights into the molecular mechanisms of the drought response in maize. This information could be used to breed drought-tolerant maize lines using molecular breeding methods.

## Materials and Methods

### Plant growth conditions and drought treatment

Seeds of maize were surface sterilized by soaking in 2% H_2_O_2_ for 10 min and rinsing with ddH_2_O. Then, the seeds were germinated in an incubator for 24 h at 28°C. Seedlings were grown in a greenhouse under a 14 h/10 h light/dark photoperiod, 60% relative humidity, and light intensity of 120 μmol m^−2^s^−1^. Seedlings were grown in half-strength modified Hoagland’s nutrient solution (pH 5.8), which was refreshed every 3 days. For the drought treatment, seedlings at the 3-fully expanded leaf stage were transferred to nutrient solution containing 20% polyethylene glycol (PEG) 6000. Leaves were harvested at 60 and 96 h of the drought treatment and after 3 d of recovery (denoted as T60, T96 and TR3d, respectively), and immediately frozen in liquid nitrogen. Control seedlings were grown under the same conditions but were not subjected to PEG treatment. Three plants from three different containers of each treatment were used as biological replicates.

### Analysis of physiological characteristics

#### Leaf relative water content analysis

The youngest fully expanded leaves were removed and weighed immediately to measure fresh weight (FW). Turgid weight (TW) was determined after leaf segments were immersed in distilled water for 6 h, and dry weight (DW) was measured after leaf segments were dried at 70°C for 24 h. Each treatment included five replicates. The relative water content (RWC) was calculated as follows: RWC (%) = [(FW–DW)/(TW–DW)] × 100.

#### Leaf gas exchange analysis

The photosynthetic rates, stomatal conductance, and transpiration rates of individual leaves were measured using a portable photosynthesis system (Li-6400; LI-COR Inc., Lincoln, NE, USA). The youngest fully expanded leaf was placed in the chamber at a photon flux density of 1000 μmol m^−2^s^−1^; the flow rate through the chamber was 500 μmol/s, and the leaf temperature was 28°C. The ambient CO_2_ concentration was approximately 380 μmol CO_2_ mol^−1^air, and the vapor pressure deficit was approximately 2.0 kPa. Five biological replicates were analyzed at the same time.

#### Osmolyte accumulation profiles

The free proline (Pro) content in fresh leaf samples (0.5 g) was determined with the ninhydrin method (Bates, Waldren, & Teare, 1973). The reaction mixture was extracted with 5 ml toluene, cooled to room temperature, and its absorbance was read at 520 nm.

#### Antioxidant enzyme activity

We determined SOD activity using an A001-1 kit (Nanjing Jiancheng Bioengineering Institute, Nanjing, China). One unit of SOD activity was defined as the amount of enzyme required for 1 mg tissue protein in a 1-ml reaction mixture to raise SOD inhibition rates to 50% at 550 nm (Tecan Infinite M200, Männedorf, Switzerland). The activity of POD was determined as described by Upadhyaya, Sankhla, Davis, Sankhla, & Smith (1985). The absorbance of the reaction mixture containing 100 ml enzyme extract, 50 mM phosphate buffer (pH 7.0), 28 ml guaiacol, and 19 ml H_2_O_2_ was read at 420 at 30 s intervals up to 2 min. The change in absorbance was used to calculate POD activity.

#### RNA extraction

Total RNA was extracted using TRIzol reagent (Invitrogen, Carlsbad, CA, USA). The RNA samples were treated with 10 units DNase I (Fermentas, Vilnius, Lithuania) for 30 min at 37°C to remove genomic DNA. The RNA concentrations were measured with a Qubit fluorometer (Invitrogen) and the RNA quality was verified using the Agilent 2100 Bioanalyzer (Agilent, Palo Alto, CA, USA) with a minimum RNA integrated number value of 8.

#### Illumina sequencing and quality controls

After total RNA was extracted, mRNA was enriched using Oligo (dT) beads. The enriched mRNA was reverse transcribed, PCR amplified, and sequenced using the Illumina HiSeq^™^ 2500 platform by the Gene Denovo Biotechnology Co. (Guangzhou, China). The total reads obtained from the sequencing machines included those containing adapters or low-quality bases, which can affect the following assembly and analysis. Thus, to obtain high quality clean reads, reads were filtered by removing those containing adapters and those containing more than 10% unknown nucleotides (N) and more than 50% low quality (Q-value≤20) bases. Last, the ribosome RNA (rRNA) mapped reads were removed through mapping reads to the rRNA database using the short reads alignment tool Bowtie2. The remaining reads were used for transcriptome assembly and analysis.

#### Relationship analysis of samples

A correlation analysis of parallel experiments evaluated both the reliability of experimental results and the operational stability. The correlation coefficient between replicates was calculated to evaluate the repeatability between samples. The closer the correlation coefficient to 1, the better the repeatability between two parallel experiments.

Principal component analysis (PCA) is a statistical procedure that converts hundreds of thousands of correlated variables (gene expression) into a set of values of linearly uncorrelated variables known as principal components. We conducted a PCA using the gmodels in the R package (http://www.r-project.org/).

#### Identification of differentially expressed genes

To identify DEGs between samples or groups, we used the edgeR package (http://www.r-project.org/). We identified genes with a -fold change ≥2 and a false discovery rate (FDR) <0.05 in a comparison as significant DEGs. The DEGs were then subjected to enrichment analysis of GO functions and KEGG pathways. The GO enrichment analysis provided all GO terms that were significantly enriched in DEGs compared with the genome background, and filtered the DEGs that corresponded to biological functions. The pathway enrichment analysis identified metabolic pathways or signal transduction pathways significantly enriched with DEGs, compared with the whole genome background.

#### miRNA sequencing and target predictions

After total RNA was extracted using TRIzol, the RNA molecules in the size range of 18–30 nt were enriched by polyacrylamide gel electrophoresis (PAGE). Then, 3’ and 5’ adapters were ligated to the RNAs, and the ligation products were reverse-transcribed by PCR amplification. The 140–160 bp PCR products were enriched to generate a cDNA library and sequenced using the Illumina HiSeq^™^ 2500 platform by the Gene Denovo Biotechnology Co. Based on the sequences of the existing miRNAs, known miRNAs, and novel miRNAs, candidate target genes were predicted using the software patmatch (v1.2) with the default parameters. The GO and KEGG pathway analyses of these genes were performed using Blast2GO.

#### Gene expression validation

We selected 24 genes with different expression patterns as revealed by RNA sequencing for validation by quantitative real-time PCR (qPCR). We extracted RNA from the leaves of three independent biological replicates for each of T0, T60, T96, and TR3d and each of their corresponding controls. First-strand cDNA was synthesized using a PrimeScript RT reagent Kit (TaKaRa, Shiga, Japan). Gene-specific primers for qPCR were designed based on the corresponding sequence using Primer5 and are listed in Supplementary Table S2 (Supporting Information). *ActinI* was used as an internal control. The qPCR analyses were carried out using SYBR Premix Ex TaqII (TaKaRa) on a LightCycler 480 instrument (Roche, Basel, Switzerland) according to the manufacturer’s instructions. Three technical replicates were analyzed for each gene.

## Results

### Physiological characteristics of *Zea mays* under drought stress and re-watering

To investigate the effect of drought stress on maize, the plants were exposed to 96 hours of water deficit, and then re-watered. Samples were collected from the drought-treated plants at 0 h, 60 h, and 96 h of drought and at 3 days after re-watering, and from well-watered (CK) plants at the same times. In CK conditions, the plants tended to grow normally with tall and straight leaves and stalks. Conversely, under drought conditions, the plants drooped. The RWC in leaves was significantly higher in the CK group than in drought-stressed group in the first 96 hours (*P* <0.01) (Fig. 1A). The differences in RWC between the drought-stressed and CK plants increased until 96 hours, then decreased after re-watering (Fig. 1A).

**Fig. 1.**
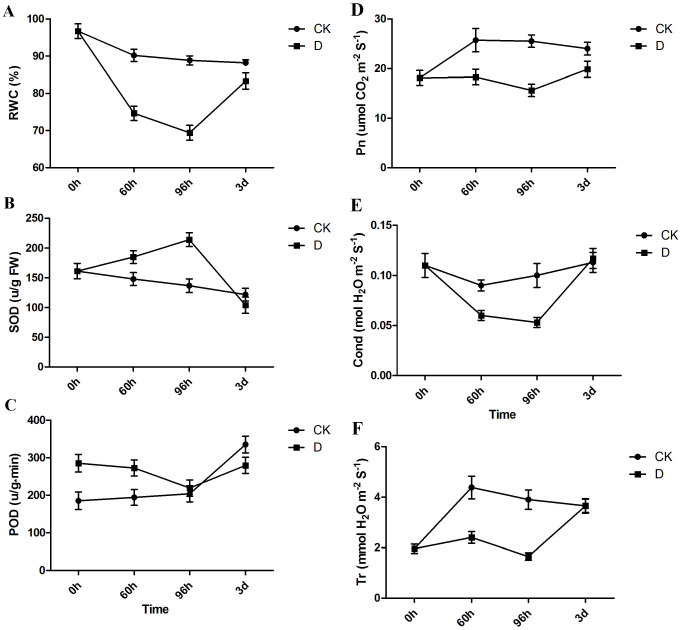
Relative water content (RWC, A), superoxide dismutase (SOD) activity (B), peroxidase (POD) activity (C), photosynthetic rate (Pn, D), stomatal conductance (Cond, E), and transpiration rate (Tr, F) of *Zea mays* during drought stress and re-watering. CK, control without drought stress; D, plants subjected to drought stress. Bars represent average (±SE) of three biological repeats.

Drought stress leads to the accumulation of reactive oxygen species (ROS), which result in cell damage. To determine the differences in ROS-scavenging capacities between the drought-stressed and CK plants, we determined the activities of POD and SOD. The SOD and POD activities were higher in the drought-stressed group than in the CK group during first 96 hours, but higher in the CK group than in the drought-stressed group after re-watering. The greatest difference in SOD values between drought-stressed and CK plants was at 96 h (Fig. 1B). After re-watering, the SOD activities were approximately equal in the CK and drought-stressed groups. The POD activity was lower in the drought-stressed group than in the CK group in the first 96 hours (Fig. 1C), but higher in the drought-stressed group than in the CK group at 3 d after re-watering.

To evaluate drought adaptation, we measured gas exchange parameters. The photosynthetic rate (Fig. 1D), stomatal conductance (Fig. 1E), and transpiration rate (Fig. 1F) decreased under prolonged drought stress, compared with their corresponding values in the control. After re-watering, the stomatal conductance and transpiration rate recovered rapidly so that their levels in the drought-stressed group were equal to those in the CK group. The photosynthetic rate in the drought-stressed also increased after re-watering, but not to the same level as that in the CK group.

### Overview of miRNA and RNA sequencing

To study the involvement of regulatory miRNAs in the complex process of the maize drought response, we profiled miRNA accumulation. After trimming adaptor sequences and removing contaminated reads, approximately 10 million tags were obtained, ranging from 16 to 35 nt in length. Of these, the 21 nt category was the most abundant, followed by the 22 nt and 24 nt categories (Supplementary Fig. S1). These were consistent with the typical lengths of plant mature small RNAs reported in other studies (Rajagopalan et al, 2006; Moxon et al, 2008; Fahlgren et al, 2007; Wang et al, 2011). In each sample, 76%–79% of the tags matched perfectly to the maize genome. Through target prediction, a total of 1028 miRNAs were predicted to regulate 11,118 genes with 22,677 target loci.

We constructed a normalized cDNA library using a mixed pool of equal amounts of 18 mRNA populations that had been extracted from leaves of plants at 60 h and 96 h of the drought treatment and at 3 days after re-watering. The library was sequenced on the Illumina HiSeq 2500 platform using the paired-end protocol. Filtering and conversion of raw reads to FASTQ format resulted in 1,201,588,344 paired-end reads with lengths of at least 2 × 150 nucleotides (Supplementary Table S1).

The resulting reads were aligned to the *Z. mays* genome that was retrieved from NCBI. The transcriptome data have been deposited to Sequence Read Archive (SRA) under the accession number PRJNA477643. The Pearson’s correlation coefficients of gene expression among repeats of each sample were relatively high. Taking CK60 as an example, the coefficient of the three repeats was greater than 0.9. The gene expression level was normalized by FPKM (Supplementary Fig. S2).

### Identification of differentially expressed genes in maize during drought and after re-watering

To identify genes with altered expression levels under drought conditions, the mRNA profiles of control plants (CK60, CK96, and CKR3d) were compared with those of plants subjected to drought stress and re-watering (T60, T96, and TR3d). Compared with the CK group in the first 60 hours, the group subjected to 60 hours of drought treatment had 3095 differentially expressed genes (DEGs), of which 1693 were up-regulated and 1402 were down-regulated. In T96 versus CK96, 942 genes were up-regulated and 999 were down-regulated. In TR3d versus CKR3d, there were significantly more down-regulated genes (4244) than up-regulated genes (1722) (Fig. 2A). In total, 478 DEGs were commonly shared among three groups (Fig. 2A). We detected 221, 226 and 215 differentially expressed miRNAs at 60 h, 96 h, and 3d, respectively, between the CK group and the drought-treated group (Fig. 2B).

**Fig. 2.**
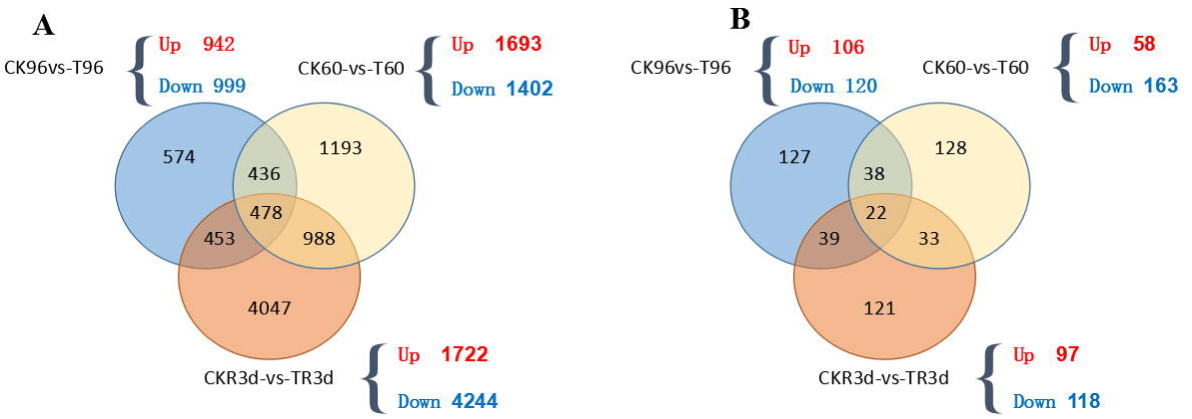
Comparisons of differentially expressed genes (DEGs) between control (CK) and drought treatments. **(A)** Venn diagram showing DEGs distribution in three libraries. (B) Venn diagram showing differentially expressed miRNA distribution in three libraries.

The miRNA sequencing results showed that in the first 60 hours of drought, 58 miRNAs were up-regulated and 163 miRNAs were down-regulated. In T96 versus CK96, 106 miRNAs were up-regulated and 120 were down-regulated. In TR3d versus CKR3d, 97 miRNAs were up-regulated and 118 were down-regulated (Fig. 2B).

According to their functional classifications, we compared the significantly regulated genes at each of the three time points (Fig. 3). On the whole, the differentially regulated genes in the first 60 hours of drought stress were related to photosynthesis and carbon metabolism, indicating a direct influence of drought on carbohydrate synthesis and metabolism (Table 1). At 96 hours, the differentially regulated genes were mainly related to phenylpropanoid biosynthesis and carbon fixation in photosynthetic organisms. After re-watering, the differentially regulated genes were mainly related to carbon metabolism and biosynthesis of amino acids. At 60 and 96 hours of drought stress, some DEGs encoded proteins involved in flavonoid biosynthesis and carbon fixation in photosynthetic organisms, indicating the common involvement of these processes in the drought stress response. After re-watering, the pathways of carbon metabolism were re-activated.

**Table 1.** Top five pathways with highest enrichment of DEGs.

| Pathway | DEGs | All genes | Qvalue | Pathway ID |
| --- | --- | --- | --- | --- |
| **CK60-vs-T60** |  |  |  |  |
| Photosynthesis | 27 | 59 | 1.01E-10 | ko00195 |
| Flavonoid biosynthesis | 10 | 23 | 1.07E-03 | ko00941 |
| Carbon metabolism | 37 | 191 | 1.09E-03 | ko01200 |
| Carbon fixation in photosynthetic organisms | 17 | 61 | 1.25E-03 | ko00710 |
| Starch and sucrose metabolism | 23 | 113 | 1.06E-02 | ko00500 |
| **CK96-vs-T96** |  |  |  |  |
| Flavonoid biosynthesis | 8 | 23 | 0.004632 | ko00941 |
| Porphyrin and chlorophyll metabolism | 9 | 37 | 0.014279 | ko00860 |
| Galactose metabolism | 9 | 38 | 0.014279 | ko00052 |
| Phenylpropanoid biosynthesis | 15 | 98 | 0.021807 | ko00940 |
| Carbon fixation in photosynthetic organisms | 11 | 61 | 0.021807 | ko00710 |
| **CKR3d-vs-TR3d** |  |  |  |  |
| Carbon metabolism | 78 | 191 | 2.57E-08 | ko01200 |
| Photosynthesis | 32 | 59 | 1.05E-06 | ko00195 |
| Carbon fixation in photosynthetic organisms | 30 | 61 | 2.93E-05 | ko00710 |
| Biosynthesis of amino acids | 65 | 191 | 3.90E-04 | ko01230 |
| Glyoxylate and dicarboxylate metabolism | 26 | 56 | 3.90E-04 | ko00630 |

**Fig. 3.**
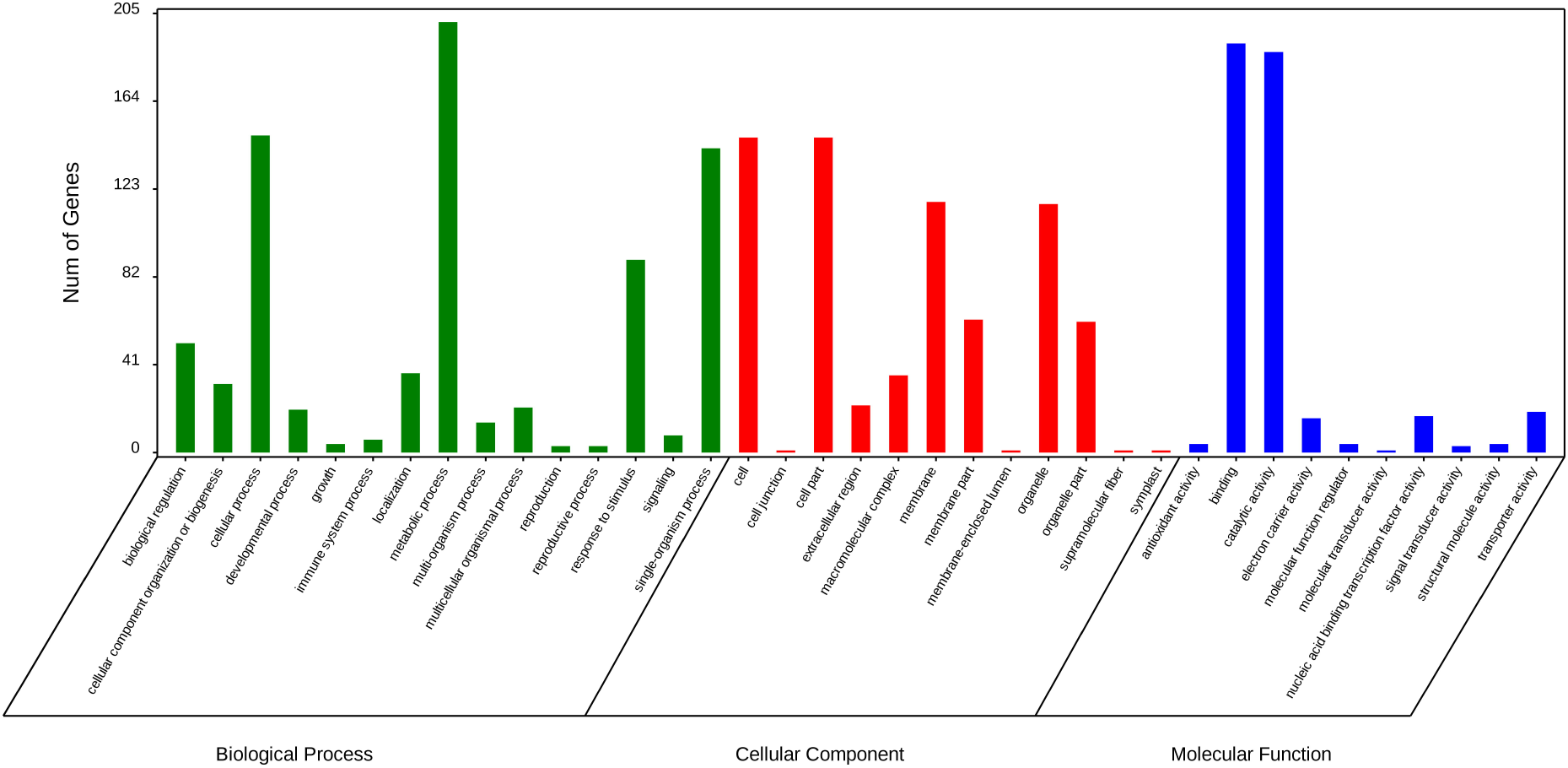
GO enrichment of common DEGs between samples.

### Identification of genes involved in cell wall

Drought caused a significant up-regulation of genes encoding proteins involved in cellular membrane stabilization and cell wall strengthening. These included the lipid-binding protein, non-specific lipid transfer protein, plant cell wall proteins, and wax synthase. Membranes are the main targets of degenerating processes that occur during drought, therefore, adaptation of membrane lipids to drought conditions might be a key mechanism for drought tolerance. Significantly more genes involved in cell wall formation were up-regulated in the first 60 hours of drought stress, but significantly more genes involved in the cell wall were down-regulated in the drought-treated group than in the CK group at 3 days after re-watering (Fig. 4A). The increased expression of cell wall-related genes may have increased the mechanical resistance of cells in drought-stressed plants. To identify the regulatory miRNAs of these DEGs involved in the cell wall, a network analysis of miRNA-genes was performed (Fig. 4B). The results showed that nine target genes were regulated by three novel miRNAs (novel-m0661-5p, novel-m0125-3p, and novel-m0283-3p) and five known miRNAs (miR9760-5p, miR5809-5p, miR5076-5p, miR6214-3p, and miR5783-5p).

**Fig. 4.**
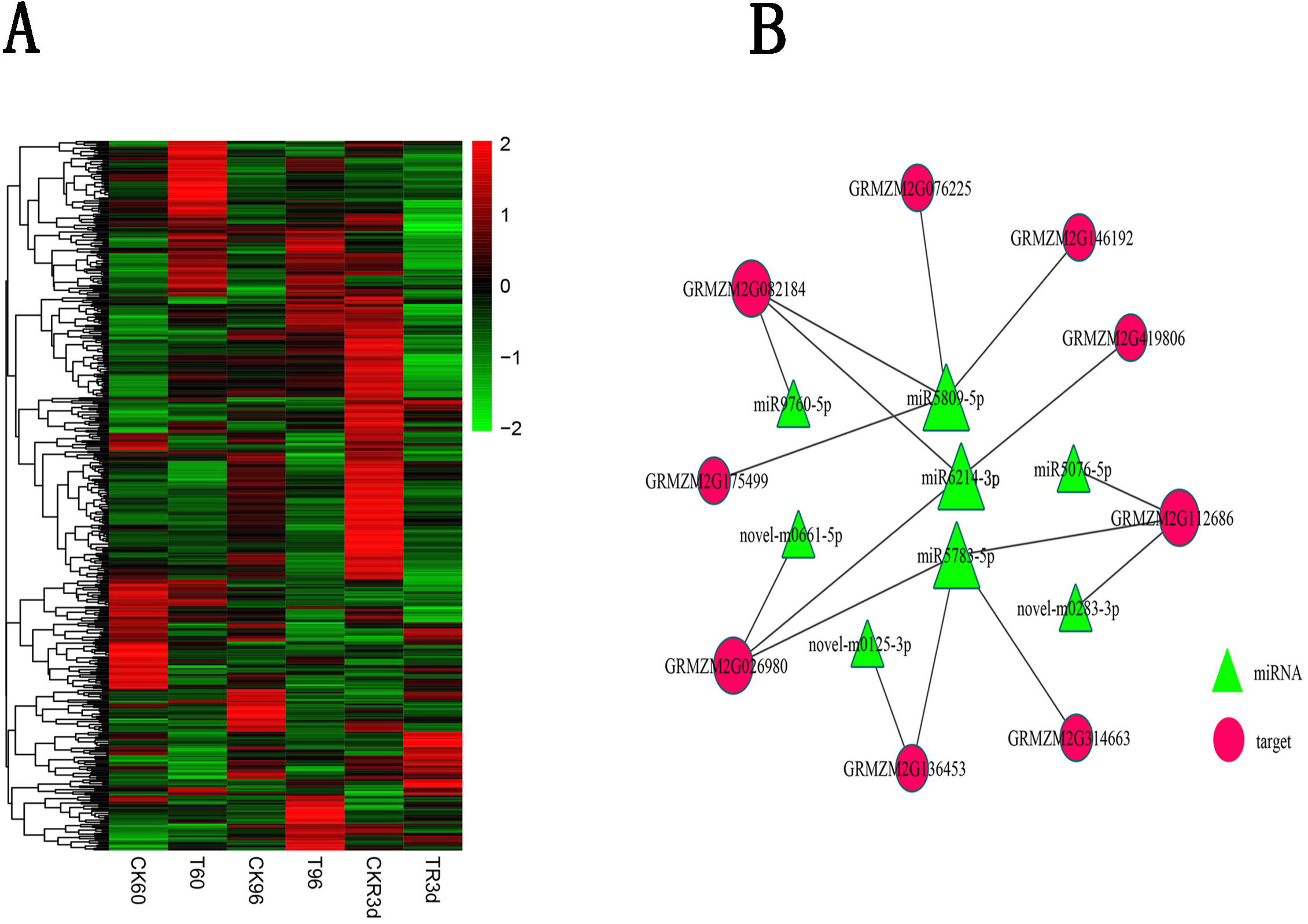
Differential expression of genes and miRNAs involved in cell wall in response to drought stress.

### Proline metabolism pathways in maize under drought stress and re-watering

Proline accumulation has long been associated with stress tolerance in plants. Proline is assumed to aid in osmotic adjustment in response to drought, however, it also plays roles in reactive oxygen species (ROS) scavenging and membrane stability. Our physiological analyzes of maize under drought stress revealed that proline significantly accumulated in maize leaves at 60 h of drought stress (*P*<0.01), but its levels did not differ significantly between drought-stressed and CK plants at other time points (Fig. 5B). We investigated the expression profiles of genes involved in proline biosynthesis in response to drought stress. We identified 26 DEGs encoding 10 enzymes involved in proline synthesis. Except for aldehyde dehydrogenase (ALDH, EC 1.2.1.3), prolyl 4-hydroxylase (P4H, EC 1.14.11.2), and spermidine synthase (EC 2.5.1.16), which were encoded by five, nine, and four genes, respectively, all the remaining enzymes were encoded by single gene (Fig. 5A). All DEGs in this pathway were clearly up-regulated in the first 96 hours of the drought treatment, except for ornithine decarboxylase (EC 4.1.1.17) and two homologs of ALDH (EC 1.2.1.3). These results suggested that maize cells increased proline synthesis in response to drought stress. The expression levels of DEGs in this pathway are shown in Fig. 5. The miRNAs analysis revealed that miR5556-5p, miR6214-3p, and novel-m0228-3p regulated three DEGs involved in proline metabolism GRMZM2G146677 (encoding aspartate aminotransferase), GRMZM2G034152 (encoding polyamine oxidase), and GRMZM2G054224 (encoding P4H), respectively.

**Fig. 5.**
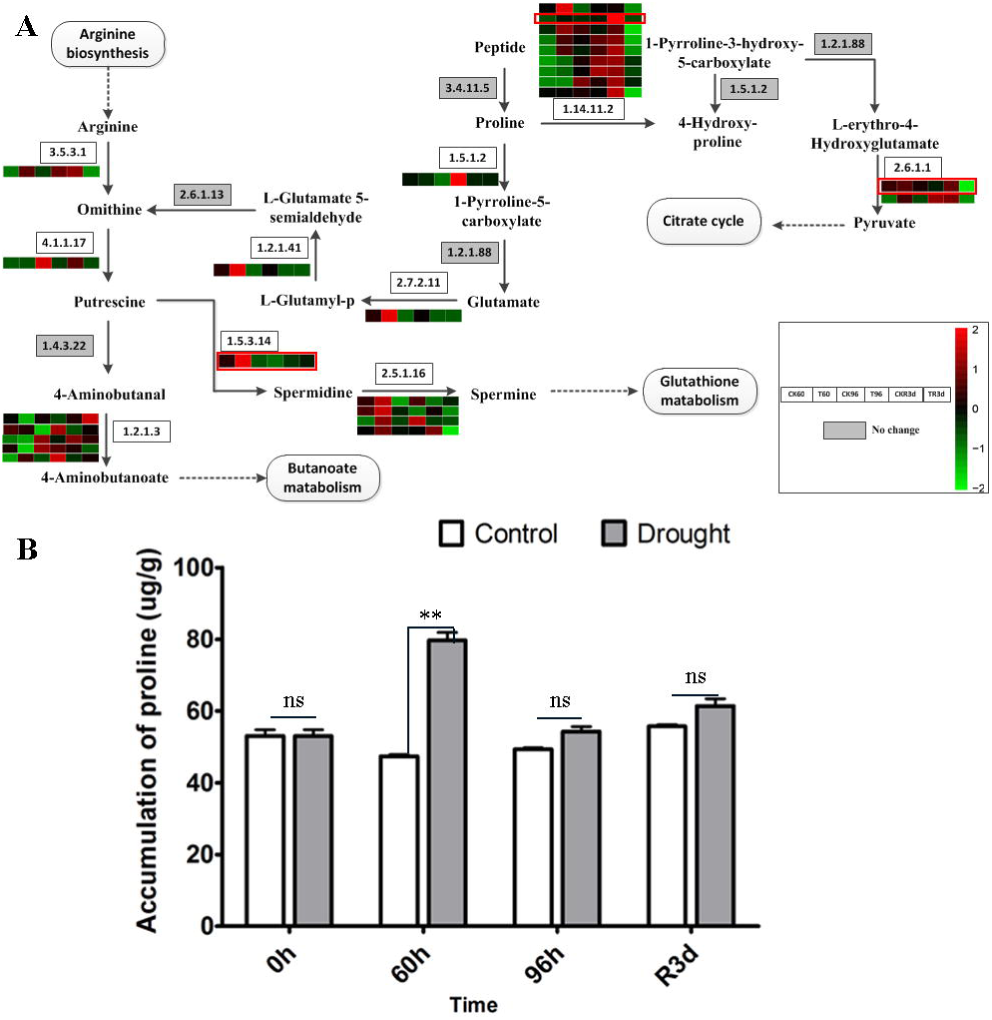
Differentially expressed genes (DEGs) involved in proline metabolism pathways, and changes in proline content in response to drought stress. (A)Expression pattern of DEGs involved in proline metabolism pathways; (B) Changes in proline content under drought stress. Genes with red box were predicted to be regulated by microRNAs. Bars represent the average (±SE) of biological repeats. Asterisks indicate statistically significant difference between groups (Student’s t-test): ^∗^*p*<0.05, ^∗∗^*p*<0.01, ns: no significant difference.

### Identification and expression analysis of transcription factors involved in the maize response to drought and re-watering

Various transcription factors and transcriptional co-activators have been found to regulate a plethora of genes and consequently confer drought tolerance. Many studies have shown that under drought stress, crops accumulate high levels of ABA, accompanied by major changes in gene expression (Galle et al. 2013; Ye et al. 2012). The ABA signaling mechanism is conserved in economically important plants, including rice and maize. Structural and computational studies have shown that the residues that comprise the gate-latch-lock components are highly conserved across species (Cao et al. 2013). High-throughput sequencing has provided information about the perception, signaling, and transportation of ABA under drought. Some of the interesting findings from crops like sugarcane are the increased abundance of bZIP factors, which may activate the transcription of drought-related genes. Here, we found that drought increased the abundance of an ABA signaling unit composed of five transcription factors (MYB/MYC, bZIP, NAC, HD-ZIP and DREB) involved in ABA-dependent or -independent gene regulation (Fig. 6A). An analysis of the miRNAs that regulate these transcription factors revealed that MYB/MYC, bZIP, NAC, and DREB were regulated by two microRNAs, while HD-ZIP was regulated by miR165-5p. The fact that MYB/MYC, NAC, and DREB were all regulated by miR5783-5p suggested that miR5783-5p plays a vital role in the regulation of the ABA signal transduction pathway in drought-stressed maize. At 60 hours of drought treatment, bZIP was activated in the drought-treated maize plants. The up-regulation of this transcription factor was related to the accumulation of dark red pigment in the drought-treated plants. The same phenomenon was observed at 96 hours of drought treatment, when bZIP was also expressed at much higher levels in drought-treated plants than in CK plants (Fig. 6B). At 3 days after re-watering, the transcript levels of the transcription factors were lower in the drought-treated plants than in the CK plants, indicating shut-down of the ABA-dependent and -independent pathways (Fig. 6A). To confirm the reliability of the DEGs identified from the RNA sequencing analysis, 24 genes involved in the ABA signal transduction pathway were selected for qRT-PCR using the specific primers listed in the Supplementary Table S1. The results of qRT-PCR were correlated with those obtained in the transcriptome analysis (Supplementary Fig. S3).

**Fig. 6.**
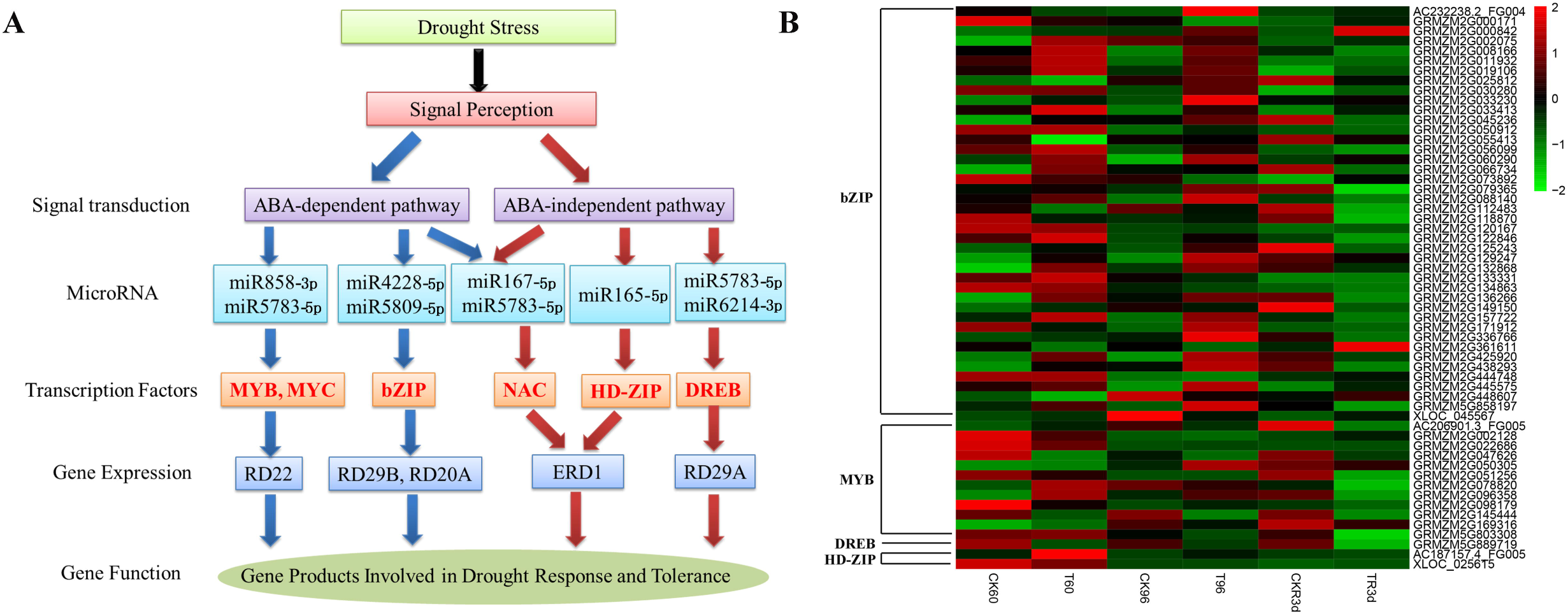
Identification and expression profile of genes involved in ABA signal transduction pathway.

### Interactions between miRNAs and their targets associated with the drought response

The results of the fluorescent quantitative analysis of miR6214-3p were consistent with the sequencing results (Fig. 9). We used the Cytoscape platform to build the network between the drought-responsive miRNAs and their targets. This allowed us to study the regulation of miR6214-3p on multiple target genes and explore its mechanism under drought stress. It was found that miR6214-3p may be involved in the regulation of the cell wall (Fig. 4B) and pro-anabolic pathways (Fig.5A), it may regulate DREB-related genes involved in ABA signaling (Fig.6A), and it may also regulate genes encoding chlorophyll a-b binding protein, as well as the BHLH and MYB transcription factors (Fig. 7). All of these genes were DEGs under drought stress (Supplementary Fig. S4), suggesting that they may play important roles in the drought response in maize.

**Fig. 7.**
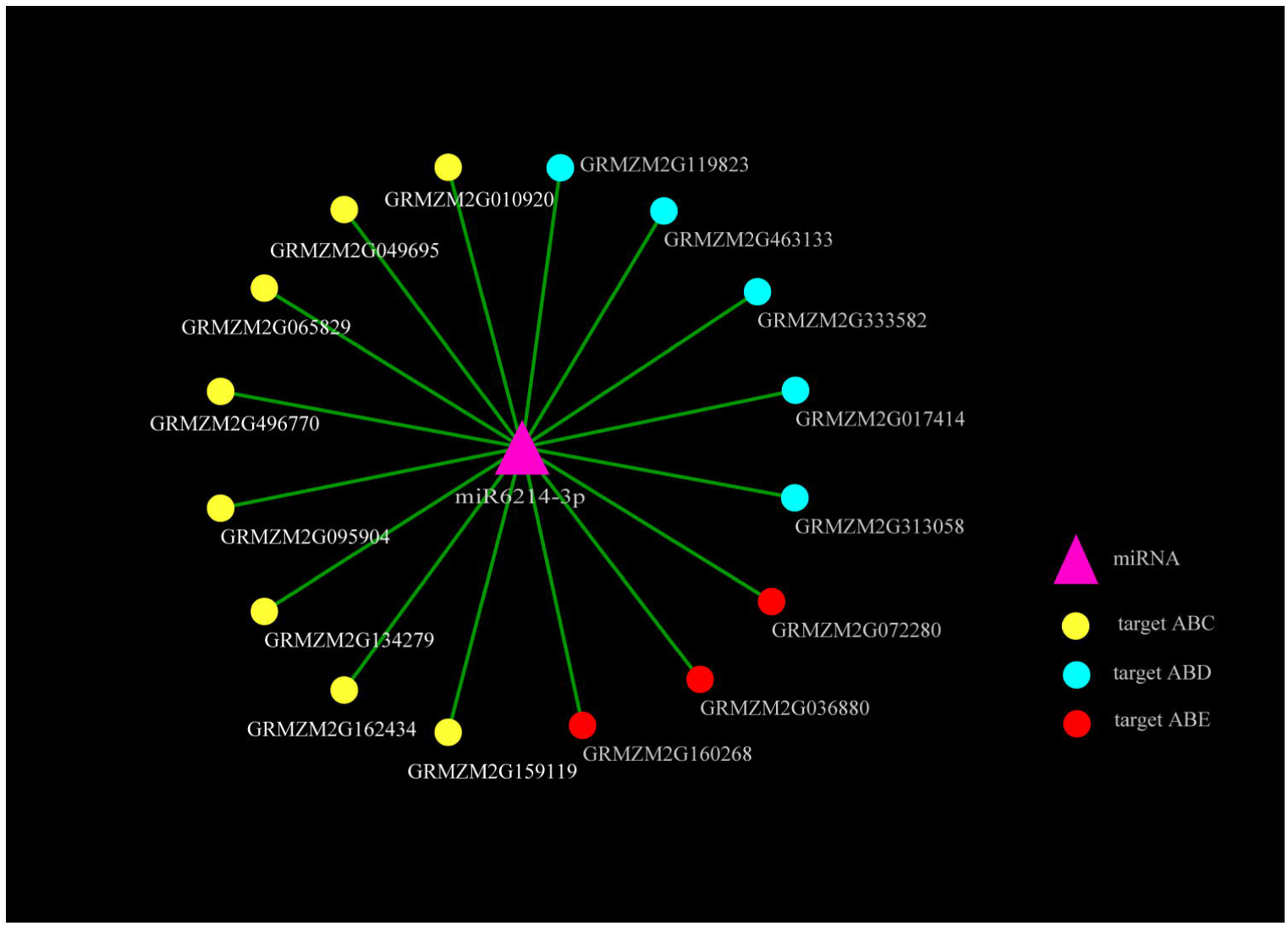
Relationships between miR6214-3p and their targets. Red circle (target ABE) indicates chlorophyll a-b binding protein; Blue (target ABD) and yellow (target ABC)circles represent BHLH and MYB transcription factors, respectively.

We selected the six miRNAs showing the largest differences in expression between drought and CK conditions, and identified their targets. Among them, up-regulated miRNAs (novel-m0414-5p) targeted eight genes associated with the drought response and tissue development (Fig. 8). These genes encoded proteins related to carotenoids (which are involved in ABA synthesis), protein phosphatase (PP2C), plant-specific serine/threonine kinase, and the ABA-responsive element binding factor. Four miRNAs were down-regulated under drought stress; Zma-miR529-5p, miR5072-3p, zma-miR167f-5p, ma-miR167j-5p, and zma-miR167e-5p. These miRNAs have potential roles in cell division, plant growth, stomatal closure, and the stress response.

**Fig. 8.**
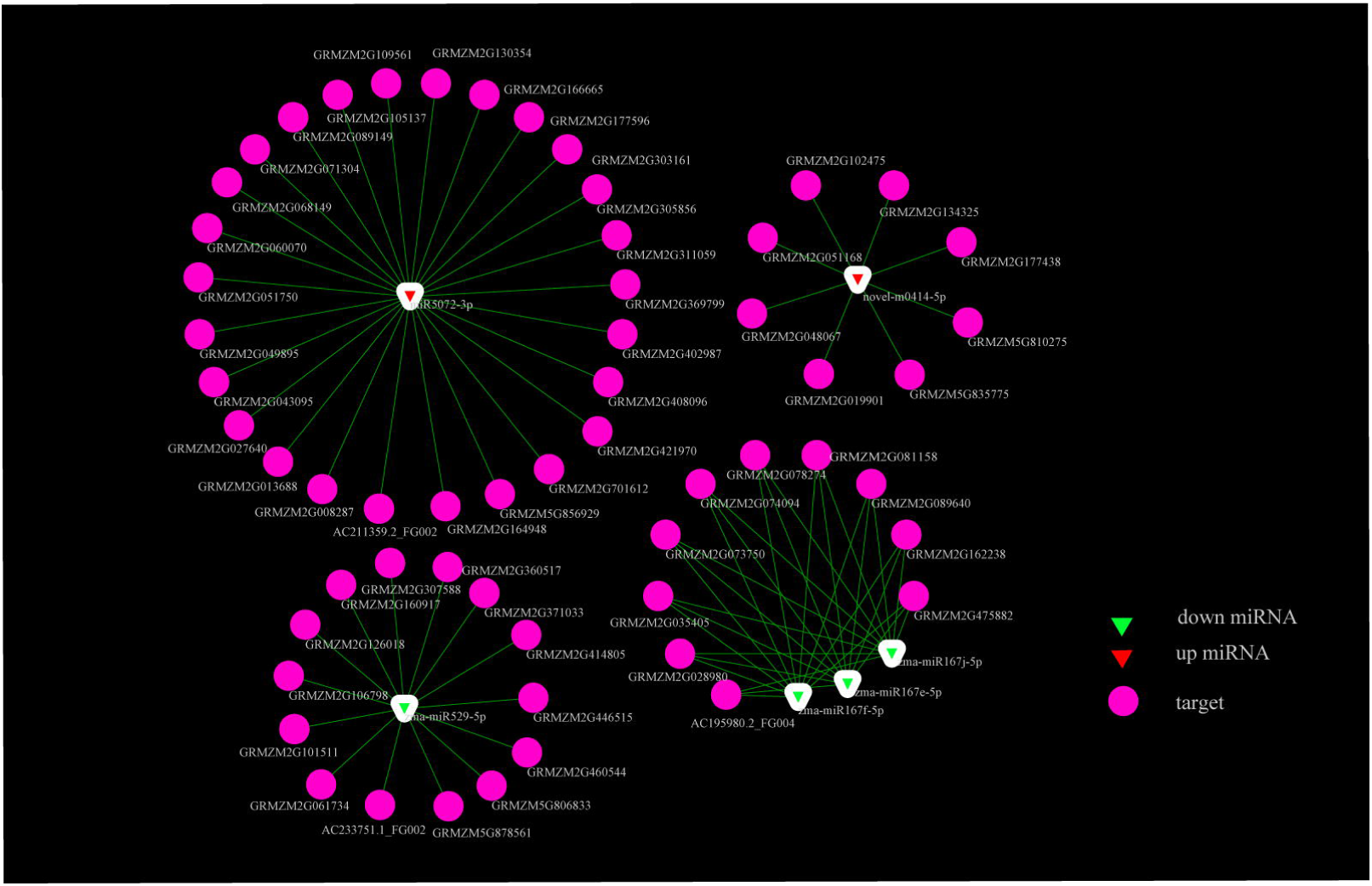
Relationships between miRNAs and their targets associated with the drought response. Red triangle indicates up-regulated miRNAs under drought stress; green triangle indicates down-regulated miRNAs under drought stress; red ellipse indicates target genes.

To confirm the reliability of miRNAs identified from the sequencing data, four key miRNAs were selected randomly for qRT-PCR using the specific primers listed in Supplementary Table S3. The results were similar to those obtained by sequencing, indicating that zma-miR167f-5p, miR5072-3p -3p, zma-miR167j-5p, and zma-miR167e-5p were down-regulated under drought stress. After re-watering, zma-miR167f-5p, miR5072-3p, zma-miR167j-5p, and zma-miR167e-5p were up-regulated (Fig. 9).

**Fig. 9.**
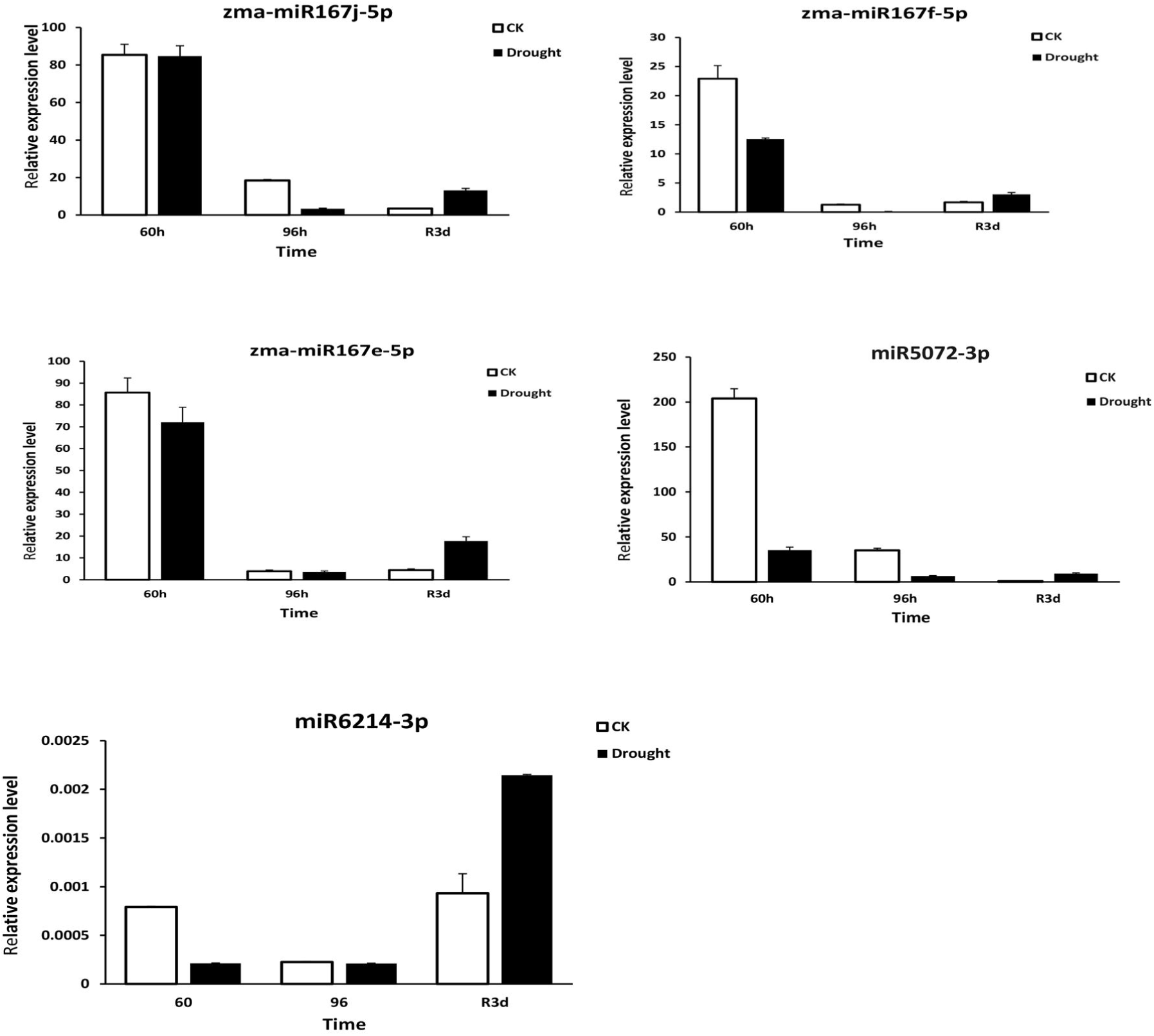
Relative expression levels of five miRNAs.

## Discussion

Drought is one of the main factors limiting plant growth and production. In practice, crops are subjected to continuous cycles of water deficit and re-watering (Perrone et al., 2012). Recent studies have shown that the capacity of plants to recover from drought is also important, particularly in crops (Luo, 2010; Perrone et al., 2012; Vanková et al., 2012; Fang and Xiong, 2015). Therefore, it is important to study the mechanisms of both drought resistance and recovery to improve crop yields under drought stress.

Drought tolerance is a complex trait. It involves multiple mechanisms that act together to avoid or tolerate periods of water deficit. Plants vary in the types and speed of responses to drought, depending on their genetic and ecotypic backgrounds. However, a number of drought responsive genes are conserved across plant taxa, especially genes encoding proteins involved in osmotic adjustment, detoxification, and cell communication and signaling. Previous studies have reported two kinds of stress-inducible genes. The first kind are genes encoding regulatory proteins (transcription factors, protein kinases, protein phosphatases) and proteins involved in signal transduction. The second kind are genes involved in water transport, cellular membrane protection, and integrity under stress conditions, scavenging of ROS (SOD, POD), and protection of macromolecules. Transcription factors that function in ABA-dependent pathways are also known to be up-regulated under drought stress (Campos, Cooper, Habben, Edmeades, & Schussler, 2004).

## Physiological responses to drought stress and re-watering

The synthesis and accumulation of ROS is promoted under stress conditions, and ROS can damage biological membranes and organic molecules. Our results showed that the activities of POD and SOD in maize leaves were significantly higher under drought stress than under CK conditions, and that their activities decreased to normal levels at 3 days after re-watering (Fig. 1B, C). This is consistent with a previous report, which noted that increased SOD and POD activities to quench ROS were related to drought tolerance in Chinese water melon (*Citrullus lanatus var. lanatus*), i.e., drought stress stimulated the enzymatic antioxidant defense system (Yan et al., 2015).

The primary reaction under drought stress is to modulate the aperture of stomata. Stomatal opening and closing modulates water loss and photosynthesis. Under drought conditions, the lack of water results in tissue dehydration in plants (Auchincloss, Easlon, Levine, Donovan, & Richards, 2014). Under such conditions, maize regulates water transport by controlling stomatal movement to prevent tissue damage. Drought can also directly affect the photosynthetic process. Drought-induced H_2_O_2_ accumulation in subsidiary cells was shown to be involved in regulatory signaling of stomatal closure in maize leaves (Yao et al., 2013). In this work, our results showed that the photosynthetic rate (Fig. 1D), stomatal conductance (Fig. 1E) and transpiration rate (Fig. 1F) were decreased by degrees under drought stress compared with CK, then recovered to normal levels after re-watering (Fig. 1 D–F).

## DEGs in maize under drought stress and re-watering

We detected 221, 226, and 215 differentially expressed miRNAs between drought-treated and CK plants at 60 h, 96 h, and 3d, respectively (Fig. 2B). Of these DEGs, 478 were commonly shared among the three groups (Fig. 2A). The GO enrichment analysis of the common DEGs among all drought-treated samples revealed the top five most enriched pathways (Table 1), which included the photosynthesis, porphyrin, and chlorophyll metabolism pathways. In response to moderate drought stress, plants modulate osmotic pressure to suppress water potential and maintain tissue turgor (Paulc, Robert, & Alvinj, 2009).

## Water stress signaling induces cellular protection processes in leaves

The cell wall is the first line of defense against abiotic stress. Many proteins that are involved in cell wall processes during normal development are also recruited during defense-related cell wall remodeling events (Maurice Bosch et al., 2011). A transcriptional analysis of GRMZ2G026980 confirmed the stimulatory effect of nitrate on xyloglucan accumulation in cells of the transition zone of the root apex; GRMZ2G026980 encodes a cell wall extension protein related to cell wall synthesis (Manoli et al., 2015; Fry et al., 1992; Geilfus et al., 2011). The expression of GRMZM2G082184 (encoding aquaporin NIP) may be associated with self-protection of plant cells under salt stress (Gao et al., 2016). The proteins encoded by three DEGs, GRMZ2G112686 (target of miR5076-5p, miR5783-5p, novel-m0283-3p), GRMZ2G026980 (xyloglucan endo-transglycosylase xyloglucan endotransglucosylase/hydrolase protein 2, target of miR6214-3p,novel-m0611-5p,miR5783-5p) and GRMZ2G082184 (aquaporin NIP, target of miR9760-5p, 6214-3p, 5809-5p) are known to participate in secondary metabolite synthesis and influence the synthesis of the cell wall. The expression levels of these genes initially decreased dramatically under drought conditions, and then increased rapidly as the drought treatment continued (Fig. 4A, B). The differential expression of these genes might be related to their involvement in the drought tolerance response.

## Effects of drought stress on proline synthesis and metabolism

Osmotic pressure is regulated by the accumulation of proline, betaine, trehalose, and fructan. The proteins encoded by GRMZM2G025867, GRMZM2G054224, GRMZM2G145061, GRMZM2G168506, GRMZM2G348578, GRMZM2G459063, GRMZM2G520535, GRMZM5G843555, and GRMZM5G855891 were all P4Hs (EC 1·14·11·2) (Fig. 5A, Supplementary Table S4). P4Hs are a large family of oxidoreductases that act on paired donors with O_2_ as the oxidant. They catalyze reactions involving the incorporation or reduction of oxygen (Bruick and McKnight, 2001; Epstein et al.,_2001). P4Hs catalyze proline synthesis and are involved in sensing hypoxia. They are upregulated in response to hypoxia, and are oxygen-dependent. Previous studies on mammals have provided clues about how these enzymes sense hypoxia. They catalyze the hydroxylation of proline to control a transcription factor [hypoxia-induced factor (HIF)] that functions as a global regulator of hypoxia in various organisms. That is, proline hydroxylation targets HIF for rapid ubiquitination and proteosomal degradation when oxygen is available (Ivan et al., 2001; Jaakkola et al., 2001).

In plants, up-regulation of ALDHs is a response to various stresses including dehydration, salinity, and oxidative stress. Plant ALDHs contribute to the synthesis of a number of osmolytes, the intracellular accumulation of which helps to counter the damaging effects of osmotic imbalance. Thus, *ALDH* gene expression appears to be a key feature of plant stress response pathways, especially those activated under oxidative stress (Zhang et al., 2012; Kotchoni et al., 2012). In this work, in addition to the novel genes XLOC_020588 and XLOC_023274, the genes GRMZM2G058675, GRMZM2G125268, GRMZM2G155502 also encoded ALDHs (EC 1.2.1.3), which play important roles in glutathione metabolism.

We determined the expression profiles of genes involved in proline biosynthesis in response to drought stress (Fig. 5 A and B). All the DEGs in the proline biosynthesis pathway were up-regulated in response to drought stress in the first 96 hours, except for ornithine decarboxylase (EC 4.1.1.17) and two ALDH homologs (Fig. 5A). These findings suggested that proline synthesis in maize was increased in response to drought stress.

## Drought stress induces ABA signaling in leaves

In plants, ABA plays a central role in many aspects of the response to various stress signals, and participates in tolerance to drought and high salinity. Previous studies have shown that the exogenous application of ABA significantly increases the ability of the plant to retain water, and that ABA accumulates to higher levels in drought-tolerant strains than in susceptible ones. In this study, many of the DEGs under drought stress and re-watering were involved in ABA metabolism. Some of the DEGs encoded transcription factors (MYB, bZIP, NAC, and DREB) involved in ABA-dependent or -independent gene regulation (Fig. 6A). The binding of ABA molecules to their receptors leads to the inhibition of PP2C, which in turn activates SNF1-related protein kinase 2 (SnRK2) (Ng et al., 2011; Soon et al., 2012). SnRK2 is an important signaling molecule that phosphorylates its downstream targets, including the transcription factors NAC, bZIP, MYB, NAC, and RAV1 (belonging to the AP2-ERF family; Furihata et al., 2006; Fujita et al., 2009; Kim et al., 2012; Feng et al., 2014). The bZIP transcription factors have been proven to play crucial roles in abiotic and biotic stress in *Arabidopsis* (Lopez-Molina et al., 2001; Uno et al., 2000; Michalak, 2006. ABA-inducible bZIP transcription factors containing ABA-responsive elements (ABRE) were shown to regulate HSFs in a drought-responsive manner (Yoshida et al., 2010; Bechtold et al., 2013). A previous report indicated that the transcription factor MYB, the target of miR858, leads to increased drought tolerance in *Ammopiptanthus mongolicus* by activating flavonoids synthesis (Gao et al., 2016). In this study, we detected an increase in the transcript levels of GRMZM2G050305, encoding the MYB transcription factor ZmMYB31, under drought stress. Its transcript levels then decreased after re-watering (Fig. 6A). ZmMYB31 is involved in the drought stress response via the ABA-dependent pathway, and its expression level is controlled by miR5809. Its up-regulation under drought conditions suggested that ZmMYB31 plays a positive role in the maize drought response. Agarwal et al. (2016) demonstrated that syntelogs of MYB31 and MYB42 bind to genes encoding enzymes in the phenylpropanoid pathway in different tissues and at different stages of development.

## miRNAs play important roles in drought stress and re-watering

It has been reported that miRNAs are involved in biological processes such as growth and development, differentiation and proliferation, and apoptosis by regulating the expression of target genes. Most research on plant miRNA has been conducted on model plants such as *Arabidopsis* and rice. Few studies have focused on the roles of maize miRNAs under stress conditions. In this study, we searched for genes related to drought resistance that are controlled by miRNAs. These results will not only help us to understand the mechanism of drought resistance in maize, but also identify potential targets to breed new drought-resistant maize varieties. In addition, the discovery of miRNAs related to drought tolerance in maize has good prospects for application. Transgenic drought-tolerant plants could be obtained by over-expressing miRNAs related to drought tolerance. This lays a theoretical foundation for the development of new drought-resistant varieties.

In a previous study, the down-regulation of miR6214-3p was associated with the activation of the stress response and antioxidant system in plants (Lu et al., 2015). In this work, we found that miR6214-3p was down-regulated under drought stress and up-regulated at 3 days after re-watering (Fig. 9). The predicted targets of miR6214-3p were involved in the metabolism and signaling pathways of the drought tolerance response (Fig. 7).

The BHLH family is one of the largest families of plant transcription factors, and its members all contain two regions (basic and HLH). Plant bHLH transcription factors bind to G-box elements in gene promoters, and are involved in growth, signal transduction, and stress tolerance. In this study, GRMZM2G313058, GRMZM2G463133, and GRMZM2G333582, which are targeted by miR6214-3p, were found to HLHs involved in drought tolerance (Fig. 7). It has been reported that the homolog of GRMZM2G313058, AtHLH112, is a transcriptional activator that mediates proline biosynthesis and increases the expression of *POD* and *SOD* genes to increase ROS scavenging and enhance stress tolerance (Liu et al., 2015). The wheat bHLH041 transcription factor, which is homolog of GRMZM2G463133 and GRMZM2G333582, was found to be associated with a high degree of cob resistance through controlling the expression of genes that strengthen the cell wall and detoxify mycotoxins (Dhokane et al., 2016).

Three predicted targets of miR6214-3p (GRMZM2G065829, GRMZM2G049695, and GRMZM2G159119) belonged to MYB superfamily, whose members are involved in signaling pathways during plant responses to biotic and abiotic stress, and regulate the synthesis of the secondary cell wall. The expression levels of these genes differed under drought conditions (Supplementary Fig. S4). MYBR35 (encoded by GRMZM2G065829) most likely functions as a transcriptional activator. Previous studies showed that DYT1 (a putative bHLH transcription factor) can activate the expression of the downstream transcription factor genes *MYB35* and *MS1* (Feller et al., 2011; Feng et al., 2012). The expression level of GRMZM2G065829 decreased dramatically during drought, then increased to higher levels than those in the CK after re-watering (Supplementary Fig. S4). The expression pattern of GRMZM2G049695 was opposite to that of GRMZM2G065829. A previous study reported that GtMYB1R1, which shares homology with the protein encoded by GRMZM2G049695, interacts with GtbHLH1 to significantly reduce anthocyanin accumulation in flowers (Nakatsuka et al., 2013). The expression level of GRMZM2G159119 decreased rapidly during drought and then increased gradually to normal levels as the drought treatment continued (Supplementary Fig. S4). This expression pattern was consistent with that reported in a previous study, where this transcription factor was shown to be involved in down-regulating genes encoding phosphoenolpyruvate carboxylase and possibly other C4-specific enzymes under nitrogen-deficient conditions (Pick et al., 2011; Urte Schlüter et al., 2012).

Among the DEGs, GRMZM2G072280, GRMZM2G160268, and GRMZM2G036880 (encoding chlorophyll a/b binding proteins) were regulated by miR6214-3p under drought stress. In another study, the down-regulation of GRMZM2G72280, which encodes the PSI component Lhca2, resulted in reduced synthesis of PSI antenna, and thus reduced light absorption under low-nitrogen stress (Mu et al., 2017). These genes were initially down-regulated under drought stress, and then recovered to normal levels as the drought treatment continued. This finding suggested that the photosynthesis of maize leaves might be inhibited under drought stress, but was able to recover as the plants adjusted their metabolic processes to adapt to the drought conditions. Consistent with this, our physiological analyses confirmed that photosynthesis recovered after re-watering.

The up-regulated miRNAs under drought was the novel-m0414-5p (Fig. 8). Meanwhile, four miRNAs were down-regulated under drought stress: zma-miR529-5p, zma-miR167f-5p, zma-miR167j-5p, zma-miR167e-5p, and miR5072-3p-3p (Fig. 8). These miRNAs have potential roles in cell division, plant growth, stomatal closure, and the stress response. Of these microRNAs, miR167 has been shown to mediate the expression of its target genes in the drought response of many plants, including *Arabidopsis*, cassava, and tomato (Kinoshita et al., 2012; Phookaew, Netrphan, Sojikul, & Narangajavana, 2014). miR5072 has been shown to modulate the synthesis of acetyl-CoA acylase by tanshinones, and the synthesis and accumulation of secondary metabolites in *Salvia miltiorrhiza*, ultimately affecting biological processes such as growth and development (Xu et al., 2014). miR529 was found to be involved in the cold response, but not in the responses to osmotic and salt stress in wheat. However, it was identified as a drought-responsive microRNA in *Oryza sativa* (Gupta, Meena, Sharma, & Sharma, 2014; Zhou et al., 2010). In our study, miR529 was down-regulated under drought stress, suggesting that it plays some role in the drought stress response in maize (Fig. 8).

In summary, drought stress resulted in decreased photosynthesis, increased antioxidant enzyme activity, and increased proline content in maize plants. Genes related to the cell wall and transcription factors regulating the ABA-dependent and ABA-independent pathways were among the DEGs under drought stress. The aim of this work was to compare changes at the physiological and gene expression levels between drought-stressed and well-watered maize plants. This allowed us identify the general responses of maize to drought stress and to explore some of the mechanisms related to the acquisition of drought tolerance.

## Acknowledgments

This research was supported by the National Natural Science Foundation of China (No. 31471452 and No. 31601258), the National Key Research and Development Program of China (No. 2017YFD0301106), the national key research and development program of china (2016YPD0101205-4) and the key grant science and technique foundation of Henan province (161100110500-0102)..We thank Jennifer Smith, Ph. D, from Liwen Bianji, Edanz Group China (www.liwenbianji.cn/ac), for editing the English text of a draft of this manuscript. We thank Guangzhou Genedenovo Biotechnology Co., Ltd for sequencing and data analyzing.

